# AP-2γ is Required for Maintenance of Pluripotent Mammary Stem Cells

**DOI:** 10.1101/2020.06.10.107078

**Authors:** Vivian W. Gu, Edward Cho, Dakota T. Thompson, Victoria C. Cassady, Nicholas Borcherding, Kelsey E. Koch, Vincent T. Wu, Allison W. Lorenzen, Mikhail V. Kulak, Trevor Williams, Weizhou Zhang, Ronald J. Weigel

## Abstract

Mammary gland ductal morphogenesis depends on the differentiation of mammary stem cells (MaSCs) into basal and luminal lineages. The AP-2γ transcription factor, encoded by *Tfap2c*, has a central role in mammary gland development but its effect in mammary lineages and specifically MaSCs is largely unknown. Herein, we utilized an inducible, conditional knockout of *Tfap2c* to elucidate the role of AP-2γ in maintenance and differentiation of MaSCs. Loss of AP-2γ in the basal epithelium profoundly altered the transcriptomes and decreased the number of cells within several clusters of mammary epithelial cells, including adult MaSCs and luminal progenitors.

AP-2γ regulated the expression of genes known to be required for mammary development including *C/EBPβ, IκBα*, and *Rspo1*. As a result, AP-2γ-deficient mice exhibited repressed mammary gland ductal outgrowth and inhibition of regenerative capacity. The findings demonstrate that AP-2γ is required for maintenance of pluripotent MaSCs and their ability to develop mammary gland structures.

**Highlights:**

- AP-2γ-deficient mice exhibited repressed ductal outgrowth and regenerative capacity
- Loss of AP-2γ reduced the number of mammary stem and luminal progenitor cells
- AP-2γ target genes, including *C/EBPβ, IκBα*, and *Rspo1*, regulate mammary development
- AP-2γ is required for maintenance of pluripotent mammary stem cells

**eTOC blurb:** Gu, Cho and colleagues utilized a conditional knockout of *Tfap2c* to examine transcriptional effects of AP-2γ on mammary stem cells. Single cell analysis demonstrated that AP-2γ-deficient mice have decreased numbers of mammary stem cells and alteration of genes required for mammary development including *C/EBPβ, IκBα*, and *Rspo1*. They demonstrate that AP-2γ is necessary for maintenance of pluripotent mammary stem cells.

## INTRODUCTION

Animal models have provided tremendous insight into the mechanisms of mammary gland development and oncogenesis with genetically engineered mouse (GEM) models allowing the identification of genes and pathways that drive these physiologic processes(McBryan and Howlin, 2017; Menezes et al., 2014). The ductal system of the mouse mammary gland develops postnatally from a single rudimentary duct that extends from the nipple during the first three weeks postpartum(Ball, 1998; Silberstein, 2001). With the start of the prepubertal period occurring at 4-5 weeks of age in mice, the mammary gland becomes responsive to reproductive hormones that initiate a growth period with terminal end bud (TEB) structures driving ductal morphogenesis. TEBs are club-shaped structures at the tip of growing ducts, which are composed of two cell types—an inner layer of body cells that give rise to the luminal mammary epithelial cells and a single layer of undifferentiated cap cells that develop into the myoepithelial layer(Morris and Stein, 2017; Williams and Daniel, 1983). The undifferentiated cap cells are thought to be pluripotential mammary stem cells (MaSCs) that are capable of giving rise to both luminal and basal progenitor cells. By 9 weeks of age, invasion of the TEBs into the stroma generates a mammary tree that fills the mammary fat pad resulting in a mammary gland characteristic of mature virgin animals.

Adult tissue stem cells are classically characterized as undifferentiated multipotent cells capable of regenerating the structure of the organ or tissue through cell division(Post and Clevers, 2019). The existence of MaSCs was first demonstrated by DeOme et al.(Deome et al., 1959) who demonstrated that normal mouse mammary tissue harvested from adult animals could reproduce the mammary gland when transplanted into the cleared fat pad of 3-week old syngeneic animals. The frequency of MaSCs defined by mammary repopulating units (MRU) in transplantation assays was estimated to be approximately 2% of basal cells(Shackleton et al., 2006; Stingl et al., 2006). More recent work has confirmed that MaSCs comprise approximately 2-5% of cells residing within the myoepithelial/basal layer of the mammary gland(Fu et al., 2019). The mouse MaSC population has the expression profile of Lin^−^ CD24^+^ CD29^high^ CD49f^high^ and also expresses the basal markers cytokeratin 5 and 14 (encoded by the *Krt5* and *Krt14* genes, respectively). Consistent with the hypothesis that pluripotent stem cells reside within the basal layer, basal cell-derived organoids are capable of forming structures comprised of inner cuboidal luminal cells capable of milk production surrounded by a myoepithelial layer with elongated morphology, whereas, luminal cell-derived organoids lack a myoepithelial layer(Jamieson et al., 2017).

Mammary gland developmental programs are characterized by changes in the transcriptome orchestrated by several transcription factors(Pal et al., 2017; Wuidart et al., 2018). The AP-2 transcription factor family is required for normal embryogenesis and the proper embryonic development of neural crest derivatives, epidermal and urogenital tissues(Hilger-Eversheim et al., 2000; Winger et al., 2006). There are five members of the AP-2 family of transcription factors—AP-2α, AP-2β, AP-2γ, AP-2δ and AP-2ε encoded by the *Tfap2a, Tfap2b, Tfap2c, Tfap2d* and *Tfap2e* genes, respectively(Eckert et al., 2005). AP-2α and AP-2γ regulate the expression of genes that define the molecular phenotype of breast cancer including estrogen receptor-alpha (ERα), HER2 and other genes predictive of outcome in luminal and HER2 breast cancer subtypes(Liu et al., 2018; Pellikainen et al., 2004; Turner et al., 1998; Woodfield et al., 2007; Wu et al., 2019). AP-2 factors also play an important role in regulating mammary gland development. In the mouse mammary gland, AP-2α and AP-2γ are expressed within the nuclei of basal and luminal mammary epithelial cells by 7 weeks of age and both factors are expressed within the body and cap cells of the TEB(Jager et al., 2010; Zhang et al., 2003). Studies have examined the effects of AP-2α overexpression driven by a transgene using the MMTV promoter(Zhang et al., 2003). The initial phase of mammary gland development was normal with mature virgin 8-week old mice expressing the transgene demonstrating normal mammary tree structures; however, at 6 months of age mice expressing the transgene demonstrated a sparser ductal network with reduced alveolar buds. During pregnancy lobuloalveolar tissue in mice expressing the transgene had reduced proliferation and increased apoptosis. Parallel studies in mice overexpressing AP-2γ demonstrated hyperproliferation, as determined by increased Ki-67 expression and BrdU incorporation, as well as increased apoptosis, with the overall effect resulting in hypoplasia of the alveolar epithelium during pregnancy(Jager et al., 2003).

Genetic knockout (KO) of *Tfap2c* results in early embryonic lethality due to loss of AP-2γ expression in extra-embryonic membranes(Auman et al., 2002). Previous studies have examined conditional knockout (CKO) of *Tfap2c* in the mammary gland. CKO of *Tfap2c* using the *Sox2* promoter to drive *Cre* recombinase showed that loss of AP-2γ impaired mammary gland branching during the prepubertal period(Jager et al., 2010). Loss of AP-2γ resulted in a reduction in the number of branch points and maximal ductal length but did not impair the generation of TEBs. By 8 months of age, the mammary ducts had completely filled the fat pad in virgin *Tfap2c* CKO mice; however, there was a reduction in the degree of branching showing that AP-2γ controls the speed of ductal elongation and the development of tertiary branches and lateral buds. In another study, CKO of *Tfap2c* using the MMTV promoter to drive *Cre* resulted in loss of AP-2γ expression in the luminal epithelial cells only(Cyr et al., 2015). Mammary gland branching was delayed in *Tfap2c* CKO mice with a reduction in branch points and an increase in the relative proportion of basal cells compared to luminal cells.

These previous studies established an important role for *Tfap2c* in mammary gland development; however, the experimental models offered a limited ability to form conclusions on the role of AP-2γ in the basal and MaSC populations. Furthermore, previous studies were not able to identify AP-2γ target genes necessary for mammary gland development. To advance our understanding of the role of AP-2γ in mammary gland development, herein we used a transgene that employs the *Krt5* promoter to drive expression of a tamoxifen-inducible *Cre-ER*, allowing control over the timing of *Tfap2c* KO and also directing the loss of AP-2γ to the basal cell population that includes MaSCs. Furthermore, performing an analysis with single cell RNA sequencing (scRNA-seq) identified AP-2γ target genes involved in mammary gland development and provided further insight into the origin and maintenance of MaSCs and progenitor cells that lead to mammary gland development.

## RESULTS

### Loss of AP-2γ Impaired Mammary Gland Ductal Outgrowth

To elucidate the role of AP-2γ in mammary gland development, we sought to examine the effect of conditional knockout (CKO) of *Tfap2c* in mouse mammary stem cells (MaSCs). MaSCs have the capacity to regenerate the mammary gland structure and demonstrate self-renewal ability(Stingl et al., 2006). Multipotent adult MaSCs reside within the basal epithelial population and express cytokeratin 5 encoded by the *Krt5* gene. To control the timing of *Tfap2c* KO in MaSCs, we utilized FVB mice expressing a tamoxifen-inducible Cre recombinase fused to the estrogen receptor with expression driven by the bovine keratin 5 promoter(Indra et al., 1999) (Figure 1a). As described for this transgene, we confirmed that expression of Cre was restricted to the cytoplasm in the basal layer with nuclear expression induced by the exposure to 4-hydroxy-tamoxifen (4OHT) (Supplemental Figure S1a). In order to minimize other genetic differences, mice carrying a floxed *Tfap2c* allele(Auman et al., 2002; Cyr et al., 2015) designed to delete exon 6 with C57BL/6 genetic background were back-crossed to the FVB/N background for 15 generations. FVB/*Tfap2c^fl/fl^/Krt5-Cre-ER^T2^* mice were pulsed with corn oil (CO) or 4OHT at a dose previously shown not to interfere with morphogenesis(Scheele et al., 2017). Mice were treated for five days beginning in early puberty at 4 weeks of age and analyzed for AP-2γ expression with immunohistochemistry (IHC) at 6, 9, and 12 weeks of age (Figure 1b & c). Significantly reduced AP-2γ expression was demonstrated in the basal cells at all time points with a slight reduction of expression in the luminal cells only at 6 weeks. Loss of exon 6 within the *Tfap2c* mRNA was detected in the basal mammary epithelial cells (MMECs) within one week of treatment with 4OHT (Supplemental Figure S1b). In parallel with examining AP-2γ expression, mammary gland development was assessed by whole mount. A delay in mammary gland ductal elongation was demonstrated with CKO of *Tfap2c* at 6, 9, and 12 weeks (Figure 2), showing that loss of AP-2γ from the adult MaSCs during this critical window profoundly affects development. However, we note that no significant differences in mammary gland ductal structure was found at 16 weeks comparing 4OHT and CO treated animals. Such an outcome might occur if recombination was either mosaic or failed to target a precursor cell population, so that wild-type cells could outcompete mutant cells and lead to an eventual recovery of ductal outgrowth. A parallel set of experiments were performed in FVB/*Krt5-Cre-ER^T2^* mice demonstrating that tamoxifen treatment using an identical protocol had no effect on mammary gland ductal morphogenesis (Supplemental Figure S2). Collectively, the findings indicate that AP-2γ plays a critical role in mammary gland ductal elongation.

**Figure 1.**
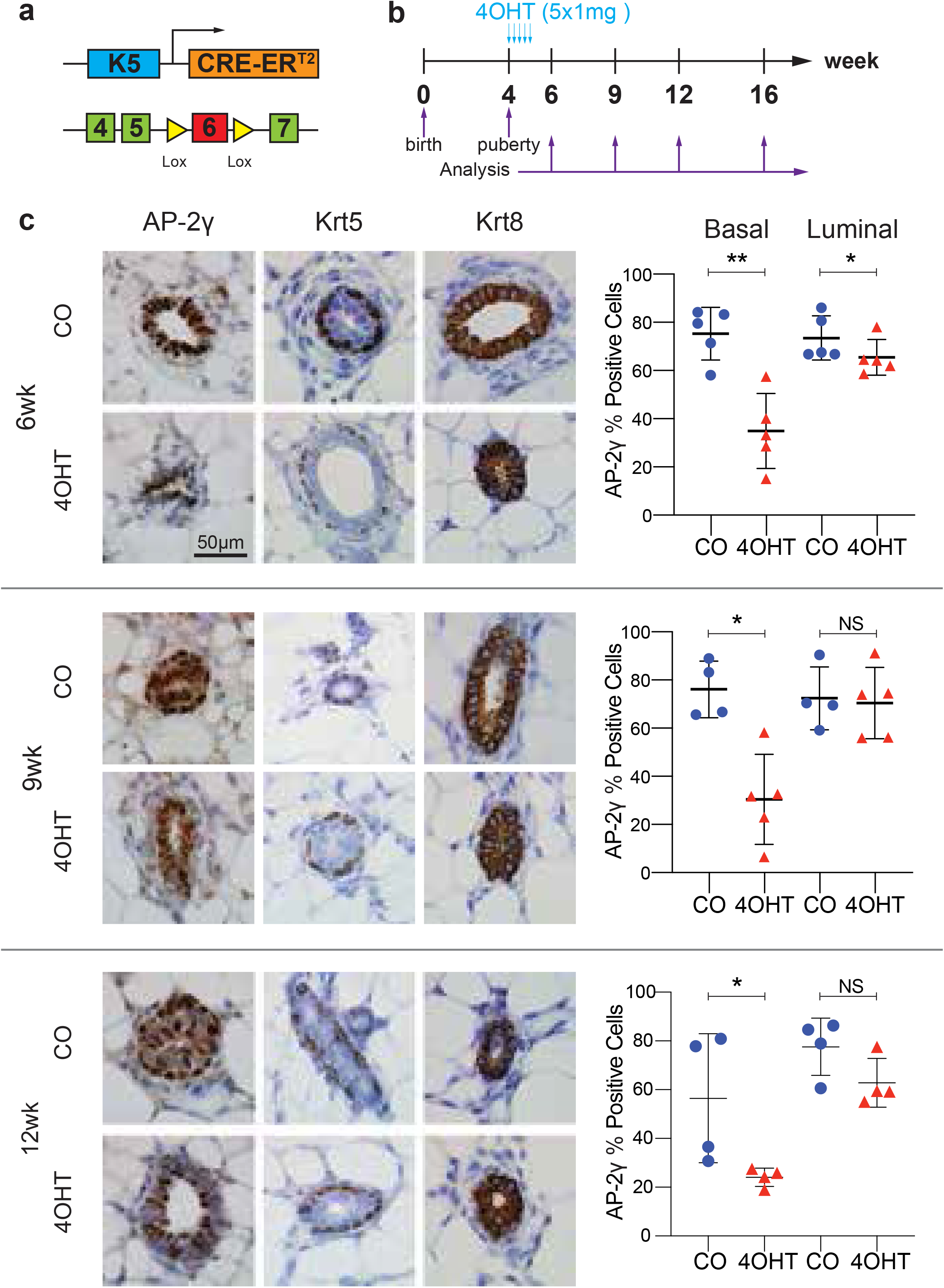
Loss of AP-2γ in basal and luminal compartments with CKO of *Tfap2*c. **a.** Krt5 promoter was used to drive expression of Cre recombinase to eliminate expression of AP-2γ through Lox sites designed to remove exon 6 of the *Tfap2c* gene. **b.** Schematic showing treatment of mice with tamoxifen (4OHT) vs. corn oil (CO) starting at 4 weeks of age for five days; analysis was performed at 6, 9, 12 and 16 weeks of age. **c.** Immunohistochemistry was used to examine loss of AP-2γ protein in MMECs with CKO of *Tfap2c*. Expression of Krt5 and Krt8 were used to demonstrate basal and luminal MMECs, respectively. There was a significant reduction of AP-2γ expression noted at 6-, 9- and 12-weeks in the basal cell compartment. Luminal cells had a significant reduction in AP-2γ expression noted at 6-weeks but this reduction did not persist at 9- and 12-weeks. *p<0.05, **p<0.01, NS: not significant

**Figure 2.**
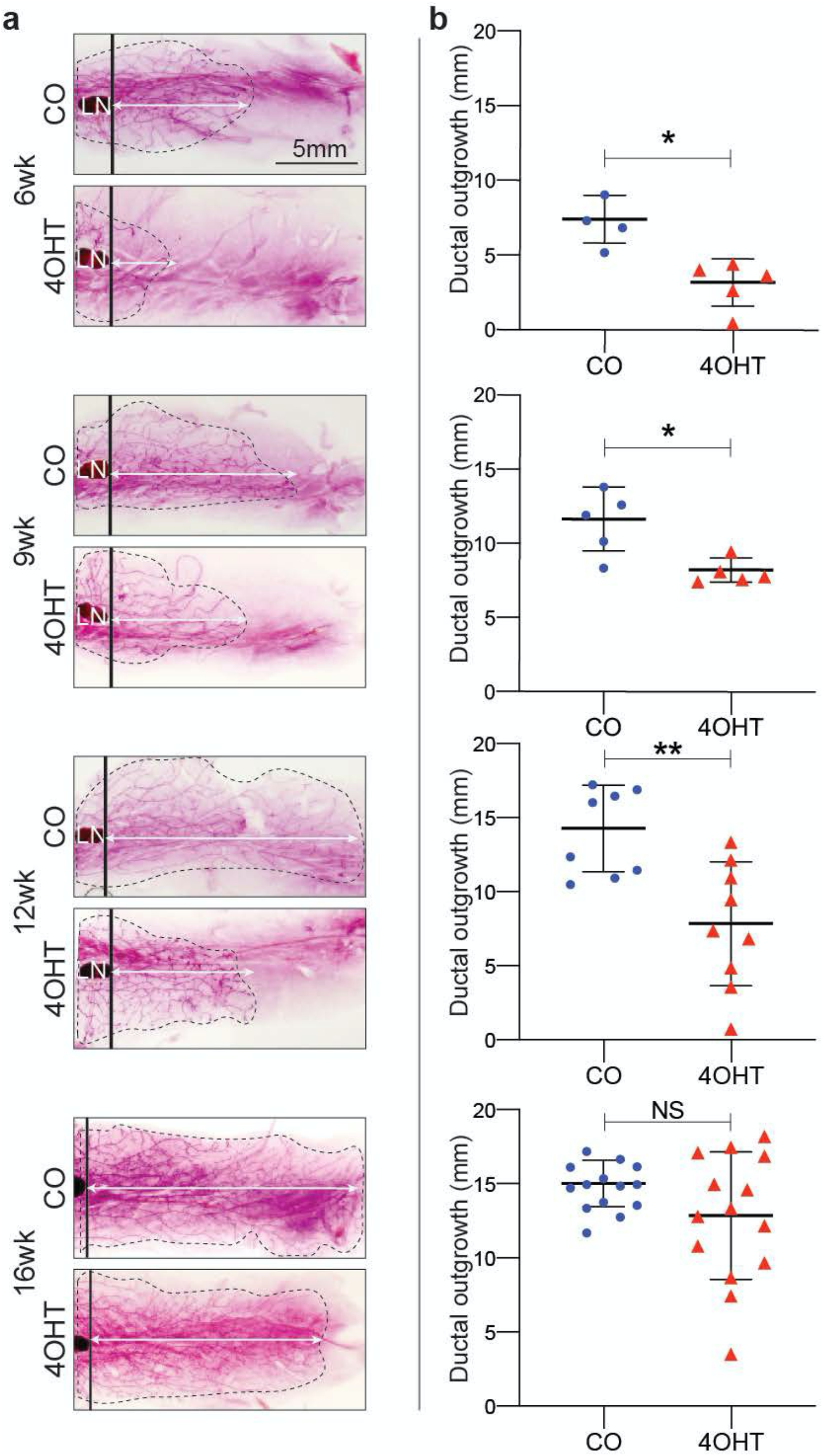
Loss of AP-2γ represses mammary gland ductal outgrowth. **a.** Representative whole mounts of mammary glands from mice treated from 4 to 5-weeks with CO or 4OHT and analyzed at 6-, 9-, 12- and 16-weeks; white arrows illustrate distances measured. **b.** Graphs showing growth of the mammary tree measured from the lymph node to most distant mammary branching demonstrating a significant reduction in growth of the mammary tree with CKO of *Tfap2c*, which resolves by 16 weeks of age. *p<0.05, **p<0.01, NS: not significant

### CKO of *Tfap2c* Alters MMEC Clusters and Patterns of Gene Expression

Having established a role for AP-2γ in mammary gland development, we sought to determine mechanistic details related to changes in gene expression within mouse mammary epithelial cells (MMECs) induced by *Tfap2c* KO. We sought to determine the extent and distribution of AP-2γ expressing MMECs and evaluate the effects and efficiency of *Tfap2c* CKO. To this end, single cell RNA-sequencing (scRNA-seq) was employed to examine changes in the transcriptome in MMECs with loss of AP-2γ. Four-week old FVB/*Tfap2c^fl/fl^/Krt5-Cre-ER^T2^* mice were similarly pulsed with 4OHT or CO and mammary glands were harvested at 9 weeks of age (Supplemental Figure S3). Single cells were generated and flow-sorted to enrich luminal and basal mammary epithelial populations, which were combined 1:1 and analyzed with scRNA-seq (Figure 3). The RNA-seq data from the combined population of 7,866 MMECs from CO-treated animals and 10,350 MMECs from 4OHT-treated animals were analyzed using uniform manifold approximation and projection (UMAP), which identified 13 distinct MMEC clusters, referred to as cluster 0-12 (Figure 3a). The pattern of expression was used to identify the major MMEC groups—luminal mature cells (LMC), luminal progenitor cells (LPC), basal cells (BC) and protein C receptor (Procr+) multipotent stem cells (PSC)(Wang et al., 2015) (Sun et al., 2018) (Figure 3b & Supplemental Figure S4). CKO of *Tfap2c* significantly reduced the relative proportion of cells within clusters 4 and 8, belonging to the basal and luminal clusters, respectively (Figure 3c, d & e). Less profound decreases were also identified in several other clusters, including cluster 10 (belonging to the BC) and 12 (belonging to the PSC). In addition, there was a significant increase in the relative proportion of cells in cluster 11, belonging to the PSC subgroup (Figure 3d & e). The *Tfap2c* gene was most highly expressed in the basal compartment with the highest percentage of *Tfap2c*-expressing cells in cluster 2 (Figure 3f and Supplemental Figure S4). Scattered cells in the luminal progenitor clusters 0, 7, 8 and 9 were also demonstrated to have relatively high *Tfap2c* expression. Although CKO repressed AP-2γ expression in the basal layer (Figure 1), we were not able to detect elimination of *Tfap2c* with 4OHT treatment from the RNA-seq data since the 3’-sequencing failed to cover the floxed exon 6 (Figure 3f).

**Figure 3.**
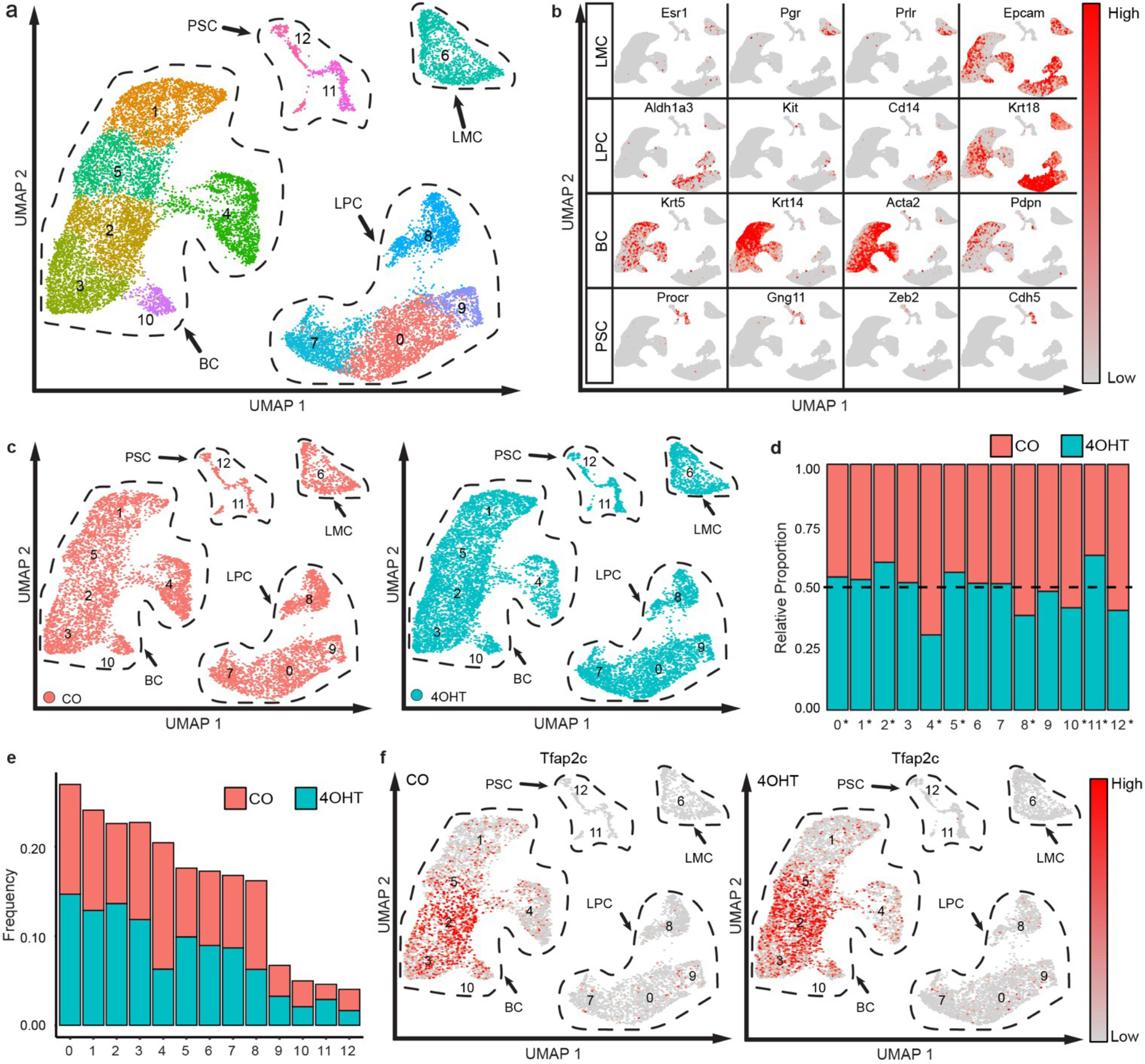
Single cell RNA-seq analysis. **a.** UMAP cluster analysis identified 13 mammary cell clusters; the transcriptome defined closely related clusters including the basal cluster (BC), luminal progenitor cluster (LPC), mature luminal cluster (LMC) and the Procr+ stem cell cluster (PSC); **b.** Examples of gene expression used to classify the major groups of mammary cell clusters; **c.** UMAP clusters shown separately for CO (orange) and 4OHT (teal) treated animals; **d.** Relative proportion of mammary cells in each cluster for CO and 4OHT treated animals, where 0.50 would indicate no change in relative number of cells; *p<0.05; **e.** Frequency of cells from CO and 4OHT-treated mice shown by the relative amount of cells by cluster; **f.** Expression pattern of *Tfap2c* in CO and 4OHT-treated mice.

The data were analyzed to identify genes with significant changes in expression with loss of AP-2γ and this gene list was subsequently subjected to Ingenuity Pathway (IP) analysis. IPA analysis indicated that many genes altered by loss of AP-2γ were involved in cell migration, invasion and differentiation (data not shown). There was a total of 644 genes that demonstrated significant changes in expression with CKO of *Tfap2c* (Supplemental Table 1). Using a difference based on a log-fold change of ±0.5 and significance with p<0.05, there was a greater number of genes repressed compared to induced with CKO of *Tfap2c* (Figure 4a). When limited to genes with an absolute percentage difference of 5% of cells in the cluster, there were 30 genes repressed and 4 genes induced by loss of AP-2γ (Figure 4b & Supplemental Figure S5). When examined by cluster, changes in gene expression were noted in every cluster, except cluster 11 (Figure 4c & Supplemental Figure S6). When considering genes that demonstrated significant changes in four or more clusters, *Cdk2ap1, Cebpb, Csn1s1, Hspa1a, Krt15, Lgals7, Plin2, Pnpla8* and *Sp140* were repressed and *Anxa3, Gadd45b, Krt18, Nfkbia, Odc1* and *Tm4sf1* were induced with CKO of *Tfap2c* (Supplemental Table 1). For all of these genes, changes in expression always included basal clusters (1, 2, 3, 4, 5 & 10), and for *Lgals7* and *Krt15*, changes in expression were restricted to the basal clusters.

**Figure 4.**
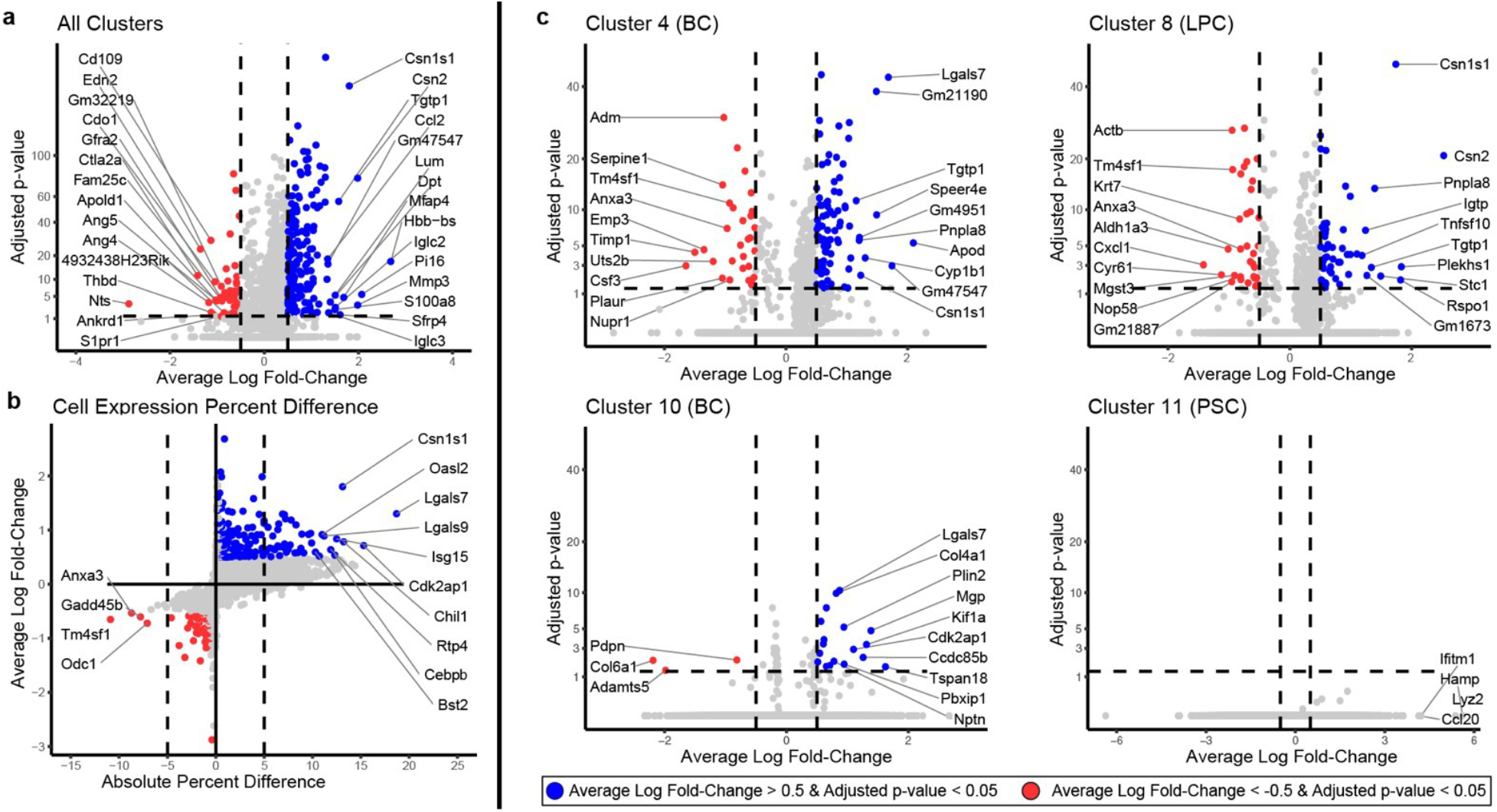
Changes in gene expression with CKO of *Tfap2c*. **a.** Volcano plot with vertical axis showing the adjusted p-value and horizontal axis average log-fold change for genes altered across all MMECs with loss of AP-2γ; colored dots represent genes with p>0.05 and average log fold change > ± 0.5; blue used for genes repressed and red used for genes induced with KO of *Tfap2c*; **b.** Same data as shown in panel **a** plotted with average log fold change versus the difference in the percentage of cells demonstrating expression of the genes. Labeled genes had an adjusted p-value < 0.05 and ± 5% percent difference; **c.** Volcano plots as presented in panel **a** for changes in gene expression for clusters 4, 8, 10 and 11.

### CKO of *Tfap2c* Repressed Pluripotency of MaSCs

Previous studies(Fu et al., 2019; Visvader and Stingl, 2014) have defined MaSCs by expression of the markers Lin^−^ Krt5^+^, CD49^high^, CD29^high^, and CD24^+^. Lgr5 has been reported as another potential MaSC marker but studies have been inconsistent showing repopulating activity in Lgr5^+^ and Lgr5^−^ subsets(Visvader and Stingl, 2014). An examination of mouse mammary glands using scRNA-seq did not identify a single, unique cluster representing MaSCs, suggesting that a functional mammary stem cell state may be generated from cells with different transcriptional signatures(Giraddi et al., 2018). Consistent with these findings, the current analysis of mammary gland transcriptome using scRNA-seq did not define a MaSC cluster specifically (Figures 3 & 5); however, based on the pattern of expression of MaSC markers, cluster 10 and a subset of cluster 4 appeared likely to represent the MaSC population. Within the basal clusters, cluster 10 had the highest percentage of cells in G1/G0 and lowest percentage of cells in S-phase (Figure 5a). CKO of *Tfap2c* reduced the number of cells within the basal clusters 4 and 10 (Figure 3d & e) and the pattern of expression was consistent with a MaSC phenotype; cells in clusters 4 and 10 were noted to be *Itga6*/CD49f^+^, *Itgb1*/CD29^High^, *Cd24a*/CD24^+^ and *Lgr5*^Low^ (Figure 5b & c). Cells with the MaSC marker expression were reduced with KO of *Tfap2c*. The impact of AP-2γ-deficiency on MaSC signature and on mammary gland ductal outgrowth suggested that CKO of *Tfap2c* reduced the frequency and/or function of multipotent MaSCs.

**Figure 5.**
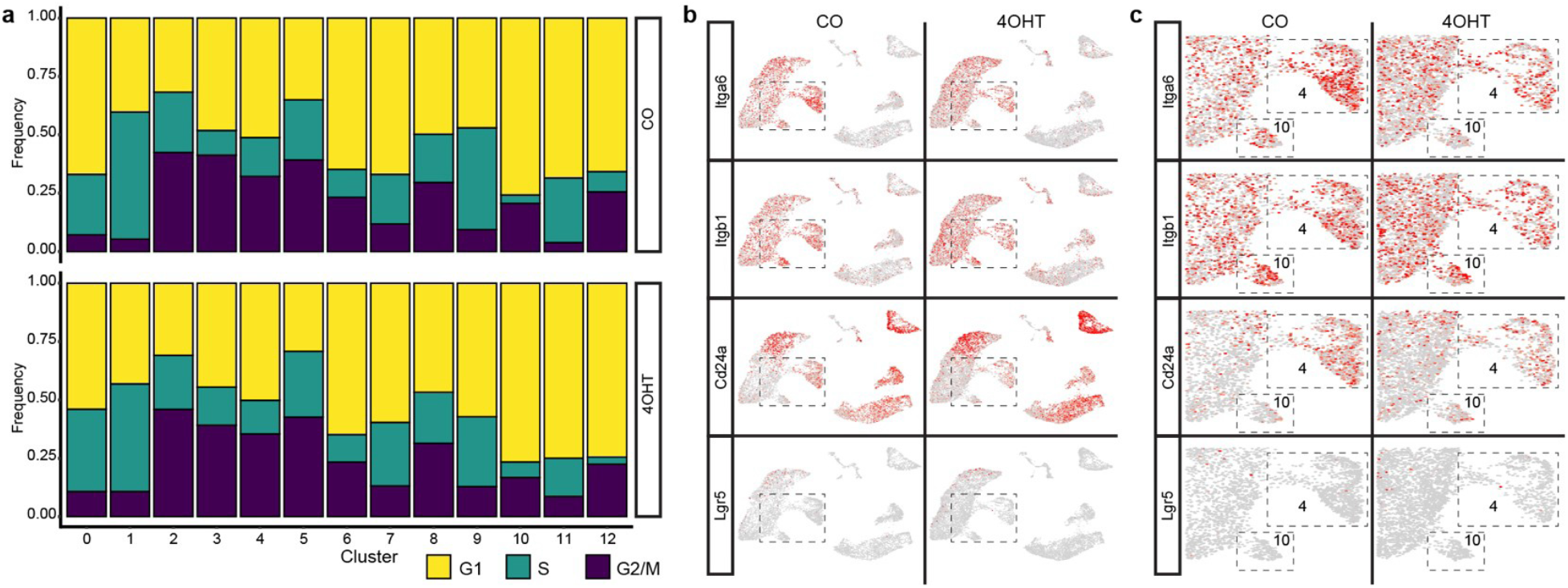
Changes in MaSCs with loss of AP-2γ. **a.** Cell cycle shown as percentage of cells in G1, S and G2/M by mammary cell cluster for CO and 4OHT-treated mice; **b.** Expression pattern for the *Itga6, Itgb1, Cd24a* and *Lgr5* genes from CO and 4OHT-treated mice; **c.** Magnified view of boxed area shown in **b** focused on basal clusters 4 and 10.

MaSCs have the ability to form mammospheres and reconstitute the mammary gland ductal structure when transplanted into the cleared fat pad of recipient mice. To examine changes in the functional properties of MaSCs, MMECs isolated from FVB/*Tfap2c^fl/fl^/Krt5-Cre-ER^T2^* mice treated with CO vs. 4OHT used in the scRNA-seq analysis were examined for the ability to form mammospheres. MMECs isolated from 4OHT treated animals demonstrated a highly significant reduction in the ability to form mammospheres, as measured by mammosphere forming efficiency (MFE), compared to CO treated mice (Figure 6a). The formation of secondary mammospheres has been advocated as a more reliable method to assess self-renewal of MaSC activity(Shaw et al., 2012). Secondary mammospheres were generated from primary mammospheres recovered from 4OHT vs. CO treated animals; primary mammospheres from 4OHT-treated mice demonstrated a reduced capacity to form secondary mammospheres (Figure 6b). Secondary mammospheres recovered from mice with CKO of Tfap2c were also noted to be smaller compared to control mammospheres.

**Figure 6.**
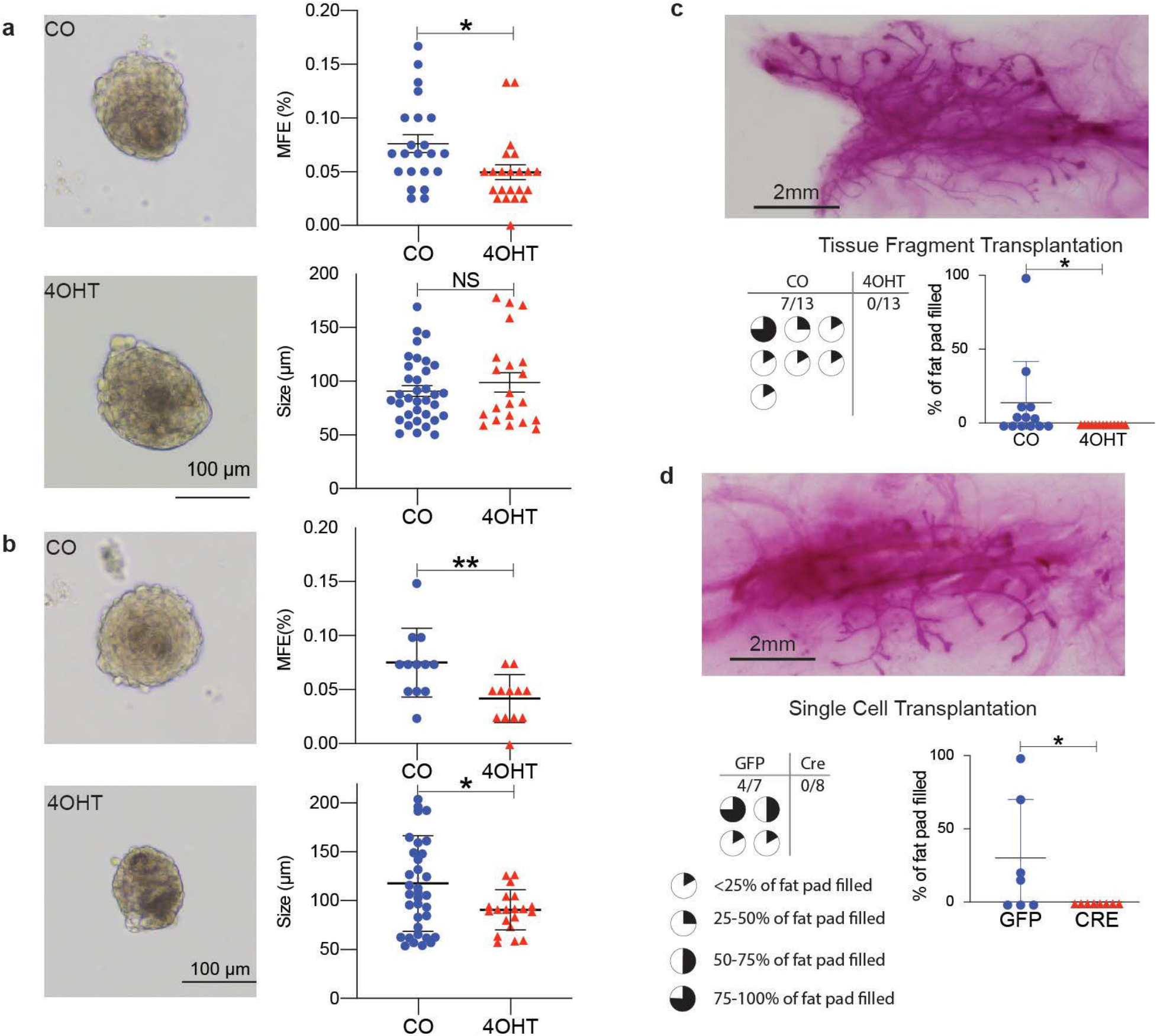
Loss of AP-2γ Reduces Mammospheres and Mammary Gland Reconstitution. **a.** Primary mammospheres formed from MMECs harvested from 9-week old FVB/*Tfap2c^fl/fl^/Krt5-Cre-ER^T2^* mice that had been treated starting at 4 weeks with 4OHT vs. CO; left panels shows examples of mammospheres and right panels show data for Mammosphere Forming Efficiency (MFE) and size of mammospheres; data shows significantly reduced MFE with KO of *Tfap2c*; *p<0.05, NS: not significant; **b.** Secondary mammospheres formed from primary mammospheres described in panel **a** shows examples of secondary mammospheres (left panels) and data demonstrating a significant reduction in MFE and mammospheres size when derived from KO animals (right panels); *p<0.05, **p<0.01, NS: not significant; **c.** Mammary tissue harvested from 9-week old FVB/*Tfap2c^fl/fl^/Krt5-Cre-ER^T2^* mice that had received 4OHT vs. CO at 4-5 weeks of age were transplanted into the cleared fat pad of 3-week old FVB mice; top panel shows examples of reconstituted mammary gland branching from CO treated animal; data demonstrated that mammary tree reconstitution was found with transplantation of mammary tissue from CO-treated but not 4OHT-treated mice; *p<0.05; **d.** Single cell MMECs from FVB/*Tfap2c^fl/fl^* mice were treated *in vitro* with Ad-GFP or Ad-Cre in parallel and injected into the cleared mammary fat pad of 3-week old FVB mice; top panel shows examples of reconstituted mammary gland branching from Ad-GFP transduced cells; data demonstrates reconstitution of the mammary tree with cells transduced with Ad-GFP but not with Ad-Cre; *p<0.05.

There is considerable debate as to whether MaSCs in the adult mammary gland are unipotent for basal and luminal lineages or whether multipotent MaSCs give rise to both basal and luminal ductal lineages during normal mammary branching(Rios et al., 2014; Van Keymeulen et al., 2011); however, there is consensus that reconstitution of the ductal tree with transplantation is a functional property of MaSCs(Visvader and Stingl, 2014). Hence, mammary gland transplantation was used to further demonstrate the effect of CKO of *Tfap2c* on functional MaSCs. Mammary tissue was isolated from 9-week old mice that had been treated with CO or 4OHT for five days starting at 4 weeks of age. Mammary tissue isolated from CO treated animals was capable of reconstituting the mammary gland when transplanted into a cleared fat pad of 3-week old syngeneic FVB mice, whereas, mammary tissue harvested from 4OHT treated animals was not able to regenerate the mammary gland structures in cleared fat pads (Figure 6c). To further examine the effect of AP-2γ on mammary gland regenerative capacity, MMEC basal cells were harvested from FVB/*Tfap2c^fl/fl^* mice and transduced with adenoviral vectors expressing Cre-GFP (Ad-Cre) or GFP alone (Ad-GFP). Basal MMECs transduced with Ad-GFP demonstrated the capacity to regenerate the mammary tree; however, cells transduced with Ad-Cre failed to demonstrate the capacity to regenerate mammary gland structures (Figure 6d). These data demonstrate a critical role for AP-2γ in maintenance of multipotent, functional MaSCs.

### Pseudotime Analysis

Pseudotime analysis of the RNA-seq data was used to generate a model of mammary gland development within the context of *Tfap2c* expression (Figure 7). The earliest MMEC precursor was represented by the Procr+ cell clusters (11 & 12). Gene expression in the Procr+ cells include expression of known mammary stem cell markers including *Zeb2, Gng11, Etv5* and *Cdh5*, but with lower expression of basal lineage marker *Krt5* (Supplemental Figure S7). The analysis is consistent with earlier studies indicating that the Procr+ cell population is a multipotent mammary stem cell population starting from the embryonic stage(Wang et al., 2015). Our data showed that Krt5^low^/Procr+ mammary cells persist in adult mammary glands during puberty but likely independent of AP-2γ (Figure 4c). Pseudotime trajectory places cluster 11 over all the branches, indicting possible microenvironmental influences on MaSC transcriptomes (Figure 7). The adult Krt5^high^/Procr-MaSCs, which are included as part of the basal cells represented as cluster 10 and a subset of cluster 4, were placed largely along the root state towards the branches of the basal and luminal divergence, likely where the basal differentiation program starts (Figure 7). The pseudotime analysis was consistent with this conclusion, showing cluster 10 and a subset of cluster 4 located adjacent and slightly after the Procr+ cluster 11. Multipotent MaSCs are capable of giving rise to basal and luminal progenitors that further differentiate into mature basal and luminal cells, respectively. Pseudotime analysis demonstrates the branching of mammary epithelial cells along the luminal or basal lineages. In the pseudotime analysis, the more terminally differentiated luminal and basal cells were seen as occupying positions at the polar ends of these differentiation pathways.

**Figure 7.**
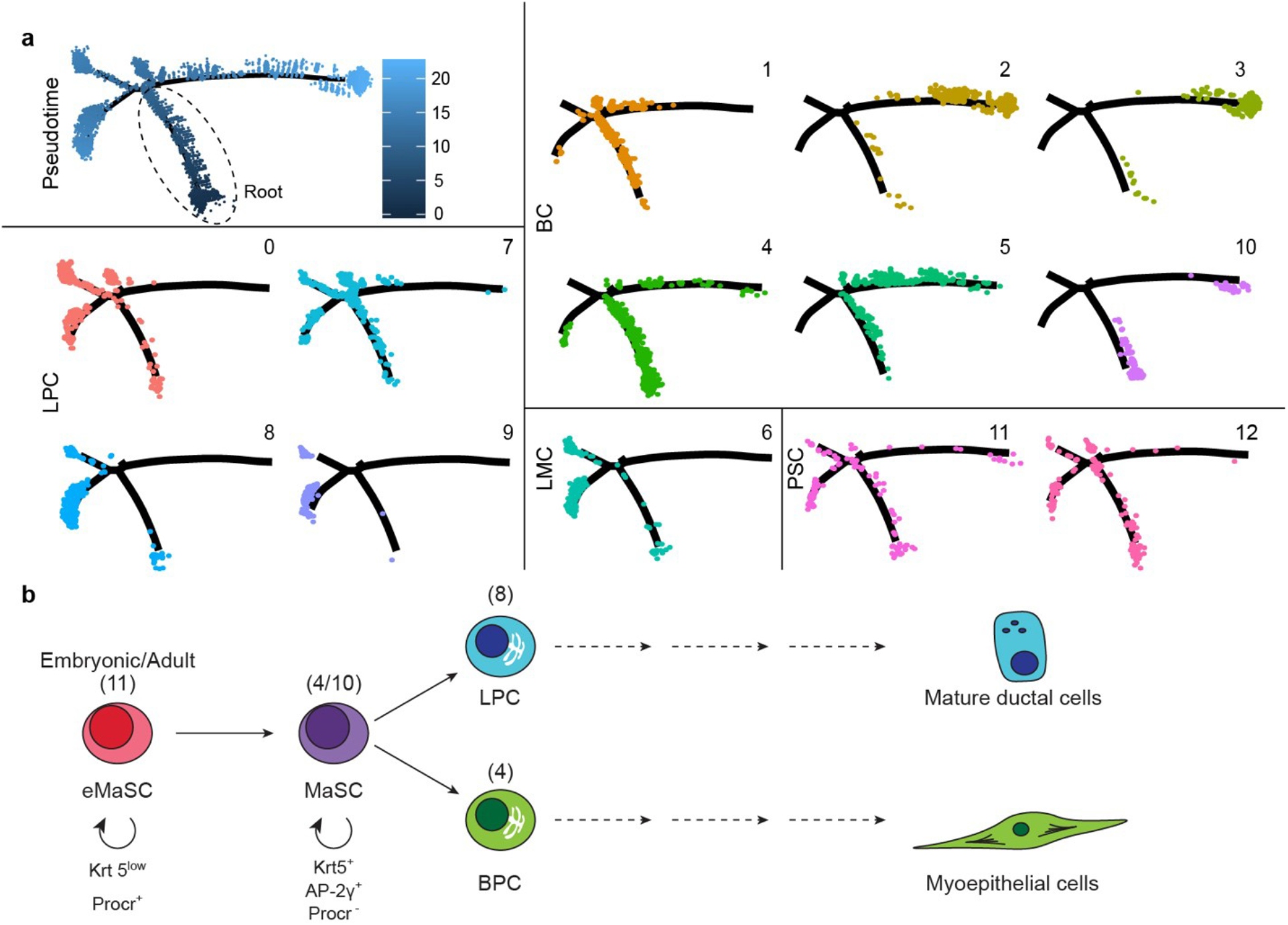
Analysis of Pseudotime and model of mammary cell development. **a.** Pseudotime analysis from RNA-seq based on data from mice treated with CO; top left panel shows relative distance from point 0 (Root, least differentiated) to more differentiated elements; remaining panels highlight location of mammary cells in each cluster organized by main cluster subtype; results show progression from the Procr+ pluripotent eMaSCs (cluster 11) to the Krt-5+ MaSC (part of cluster 4 and cluster 10), basal progenitors, luminal progenitors, mature basal and mature luminal cells; **b.** Model of mammary development supported by results with CKO of *Tfap2c*; the results indicate that KO of *Tfap2c* in pluripotent Krt5+ MaSCs blocks the multipotent ability of MaSCs.

## DISCUSSION

The current study provides important insight into the role of AP-2γ during the prepubertal stage of mammary gland development. CKO of *Tfap2c* within the Krt5+ basal cell compartment repressed mammary gland ductal outgrowth and inhibited the ability of mammary epithelial cells to form mammospheres and regenerate mammary gland structures. The phenotypic alterations with loss of AP-2γ expression were accompanied by changes in gene expression within the basal cells and also included significant alterations in gene expression in luminal cell clusters. Where they can be compared, the current findings are consistent with previous reports that examined the role of AP-2γ in the mouse mammary gland(Cyr et al., 2015; Jager et al., 2010). As noted previously, *Igfbp5* was identified as an AP-2γ target gene in the mammary gland(Jager et al., 2003) and we noted a significant repression of *Igfbp5* expression in basal cluster 4 and luminal cluster 7 (Supplemental Table 1). The current findings extend previous reports by clarifying the role of AP-2γ in the basal/MaSC compartment and by identifying AP-2γ target genes that regulate mammary gland development.

There is considerable debate concerning the potential role of multipotent Krt5^+^ MaSCs within the basal compartment in mammary gland outgrowth during the pubertal period. Lineage-tracing experiments using Krt5-CreER/Rosa-YFP mice treated with tamoxifen at 4 weeks of age demonstrated labeling of the basal compartment exclusively(Van Keymeulen et al., 2011). Similar experiments using a Krt14 or Krt8 promoter to drive Cre expression yielded similar findings with exclusive labeling of either the basal or luminal compartment, respectively. On the other hand, transplantation experiments demonstrated that CD29^high^CD24^+^ basal cells were capable of generating normal mammary gland structures. The results supported the conclusion that during pubertal development, distinct unipotent MaSCs give rise to the basal and luminal cell compartments; however, mammary gland transplantation disrupts the normal relationship of basal and luminal cells uncovering the reconstitution capacity of Krt5^+^ MaSCs. In contrast to this study, lineage-tracing experiments reported by Rios et al.(Rios et al., 2014) using high resolution 3D imaging demonstrated labelling of both the luminal and basal populations of the ductal tree when driven by the *Krt5* promoter, supporting the model of a multipotent Krt5^+^ MaSC giving rise to mature basal and luminal cells during pubertal mammary gland development. The existence of multipotent Krt5^+^ MaSCs supports their role in mammary gland development during puberty, and would not be inconsistent with the presence of unipotent basal and luminal progenitor cells in adult mammary glands. It has been suggested that the model employed by Van Keymeulen et al. may have predominantly labelled basal progenitors(Visvader and Stingl, 2014).

Our findings further support a role for multipotent Krt5^+^ MaSCs. The current results identified changes in gene expression in both basal and luminal cell clusters. The most likely basis for this finding is that CKO of *Tfap2c* driven by the *Krt5* promoter eliminated AP-2γ expression in multipotent MaSCs that eventually gave rise to both basal and luminal progenitors with KO of *Tfap2c*. A potential conundrum is that the data demonstrate that loss of AP-2γ inhibited the multipotent function of MaSCs evidenced by a decrease in the ability to form mammospheres and reconstitute mammary gland structures. It seems likely that expression of Cre resulted in *Tfap2c* gene disruption but that the AP-2γ protein persisted for a time allowing multipotent MaSCs to divide giving rise to basal and luminal progenitors, which carried the deleted *Tfap2c* gene. Previous studies in luminal breast cancer showed that knockdown of *TFAP2C* induced epithelial-mesenchymal transition with repression of luminal gene expression and induction of basal gene expression(Bogachek et al., 2016; Cyr et al., 2015). One likely hypothesis is that AP-2γ is necessary for the multipotent MaSCs to generate luminal progenitors, which could explain both the findings in mammary gland development and luminal breast cancer models (Figure 7b). Further evidence supporting this hypothesis is the finding that early after CKO of *Tfap2c* reduced expression of AP-2γ was identified in the luminal compartment but significant reduction in AP-2γ was not identified at later time points (Figure 2). If CKO of *Tfap2c* in MaSCs blocked their ability to give rise to luminal progenitors, then all subsequent luminal epithelial cells would have to be derived from MaSCs with an intact *Tfap2c* gene, and would explain why decreased AP-2γ expression is not detected at later time points.

It was particularly interesting that Cluster 11, composed of Procr+, Krt5-cells, was the only cluster without significant changes in the pattern of gene expression with CKO of *Tfap2c*. This finding suggested that cluster 11 represents an early multipotent MaSC population, likely the embryonic MaSCs (eMaSCs) due to its expression at embryonic stages of mammary placode(Wang et al., 2015) that gives rise to the Krt5+ MaSCs within the basal cell population (Figure 7b). Our data is consistent with the findings of Wang et al.(Wang et al., 2015), which identified the Procr^+^ Krt5^low^ Krt14^low^ mammary cells as a multipotent eMaSCs at the top of the developmental hierarchy capable of giving rise to Krt5^+^ MaSCs. The lack of changes in expression of AP-2γ target genes in cluster 11 is consistent with the hypothesis that the Procr+ cells in cluster 11 precede expression of *Krt5* and therefore have an intact *Tfap2c* gene.

Interestingly, although several analyses of mammary gland using scRNA-seq have been reported, this is the first scRNA-seq data that confirmed the existence of Procr+ MaSC population represented within clusters 11/12. Although pseudotime trajectory analysis demonstrated the bulk of these cells at earliest developmental stage, they are also found scattered throughout the pseudotime branches, likely due to either the microenvironmental influence at particular locations or the heterogeneity of their transcriptome signature. This broader mammary distribution of MaSCs has been discussed before(Pal et al., 2017). This likely represents the population of MaSCs spread within the entire mammary gland throughout mammary development processes that have very low cycling cells based on cell cycle analysis(Wang et al., 2015) (Figure 5a). Upon injury, these are likely the cells leading to mammary repair at the injured location. AP-2γ-responsive Krt5+ populations, represented within basal clusters 10/4, include a subset of Krt5+ MaSCs, which are likely the cap cells located within the terminal end buds during pubertal development of the mammary gland. These clusters represent a highly proliferative stem cell population that are heterogeneous in their transcriptome profile(Scheele et al., 2017). It is worth noting that the decrease in AP-2γ-responsive Krt5+ populations in clusters 4/10 likely triggers a compensatory signal to Procr+, Krt5^Low^ eMaSC, which leads to the compensatory increase in Procr+ MaSC in cluster 11. The data suggest that following the pulse with tamoxifen, the compensatory increase of Procr+ cells divide to generate Krt5+ MaSCs with an intact *Tfap2c* gene, which lead to mature basal and luminal ductal cells resulting in normal mammary morphogenesis by 16 weeks of age. However, these data do not address whether multipotent Procr+ cells can lead directly to luminal cell types independent of the Krt5+ MaSC (Figure 7b).

Several genes known to regulate mammary gland development were altered by CKO of *Tfap2c* including *Cebpb, Nfkbia* and *Rspo1*. The *Cebpb* gene (also known as *C/EBPβ)* was significantly repressed in several basal and luminal mammary epithelial cell clusters with loss of AP-2γ. *C/EBPβ* encodes three isoforms of a b-ZIP transcription factor translated from different translational start sites, referred to as LAP1, LAP2 and LIP(Zahnow, 2009). Mice deficient for *C/EBPβ* demonstrate disrupted mammary gland morphogenesis with reduced growth and secondary branching and loss of secretory cells during pregnancy(Robinson et al., 1998; Seagroves et al., 1998). *C/EBPβ* - deficient mice also lack expression of β-casein, and the *Csn2* gene (encoding β-casein) was significantly repressed in luminal clusters 0, 7 and 8 with CKO of *Tfap2c* (Supplemental Table 1). Of note, the *Csn1s1* and *Csn1s2a* genes (encoding casein-αS1 and casein alpha s2-like A, respectively) were similarly repressed by CKO of *Tfap2c* in basal and luminal cell clusters. Mice overexpressing the *C/EBPβ* LIP protein isoform in the mammary gland develop alveolar hyperplasia, further supporting a critical role for *C/EBPβ* in regulating mammary gland development(Zahnow et al., 2001). IκBα encoded by the *Nfkbia* gene is a major inhibitor of the NFκB transcription factor(Henkel et al., 1993). Mammary epithelial cells deficient in IκBα when transplanted into syngeneic animals develop an increase in lateral ductal branching and intraductal hyperplasia(Brantley et al., 2001). This finding indicates that NF-κB is a positive regulator of mammary epithelial proliferation; since CKO of *Tfap2c* increased expression of *Nfkbia* in several basal clusters, the findings are consistent with loss of AP-2γ inhibiting mammary gland development through repression of NF-κB. Loss of *Rspo1* expression resulted in an absence of mammary side-branching and loss of alveolar development during pregnancy(Chadi et al., 2009) and CKO of *Tfap2c* resulted in repression of *Rspo1* expression in several of the luminal epithelial cell clusters. Taken together our findings indicate that loss of AP-2γ in the MaSC compartment leads to repression of mammary gland branching and normal alveolar development, at least in part, through the regulation of several drivers of mammary proliferation, branching and alveolar maturation including *C/EBPβ, IκBα*, and *Rspo1*.

Many of the genes regulated by AP-2γ in MMECs are known to be involved in cell migration, invasion, differentiation and breast cancer tumor growth. The expression of *Cdk2ap1* was significantly repressed in nearly every one of the cell clusters. The Cdk2ap1 protein binds to and inhibits the activity of CDK2 resulting in altered proliferation and cell cycle regulation(Kim et al., 2005; Shintani et al., 2000). CDK2AP1 regulates the expression of the pluripotency genes OCT4 and NANOG and represses differentiation of embryonic stem cells through regulation of pRb, though its effect on gene regulation and differentiation appears to be cell-type specific(Alsayegh et al., 2018; Kim et al., 2005). Interestingly, *Rb1* was repressed in cluster 0 (the largest luminal cell cluster) with CKO of *Tfap2c*. In breast cancer, CDK2AP1 acts as a tumor suppressor as demonstrated by its ability to reduce proliferation, inhibit colony formation and represses *in vivo* tumor growth(Zhou et al., 2012). The expression of *Anxa3* was induced in several basal and luminal clusters with CKO of *Tfap2c*. Silencing *ANXA3* expression in human breast cancer cells increased the percentage of cells in G0/1 and the rate of apoptosis with reduced proliferation, migration and invasiveness(Li et al., 2018; Zhou et al., 2017). The expression of *Lgals7* (encoding galectin-7) was repressed in all basal clusters with CKO of *Tfap2c*. Although *Lgals7*-deficient mice develop normal mammary glands, these mice demonstrate a delay of mammary tumor growth when expressing the MMTV-*Neu* transgene(Grosset et al., 2016). There is also evidence that galectin-7 may participate in oncogenesis induced by mutant p53 in breast cancer(Campion et al., 2013). Loss of AP-2γ induced expression of *Cxcl1* in basal and luminal clusters. The *Cxcl1* gene encodes chemokine (C-X-C motif) ligand 1, which is a member of the chemokine family involved in inflammation and tumor progression(Dhawan and Richmond, 2002). Cxcl1 is up-regulated in ERα-negative breast cancer and stimulates migration and tumor invasiveness through Erk1/2 activation(Yang et al., 2019). Breast cancers with overexpression of CXCL1 have increased propensity for survival at metastatic sites and demonstrate chemoresistance(Acharyya et al., 2012).

In summary, AP-2γ plays an important role in normal mammary gland development and ductal outgrowth during the prepubertal period. Loss of AP-2γ reduced the number of functional MaSCs and significantly repressed the capability for MaSCs to regenerate mammary gland structures. The data suggest that loss of AP-2γ inhibits the ability for multipotent MaSCs to differentiate along the luminal lineage. Changes in the pattern of gene expression with CKO of *Tfap2c* indicate that a Krt5^Low^, Procr+ cell population gives rise to Krt5+ MaSCs that are capable of generating differentiated basal and luminal cells in the mammary gland. Within the basal and luminal MMECs, AP-2γ regulates the expression of several genes known to be necessary for mammary gland development including *C/EBPβ, IκBα*, and *Rspo1*. Additional genes involved in regulation of proliferation, invasiveness and differentiation were found to be AP-2γ target genes in the mammary gland. The model system described will further allow an examination of AP-2γ influence of MaSCs during other stages of mammary gland development and lactation and will provide an important resource to further investigate the role of AP-2γ in mammary oncogenesis.

## EXPERIMENTAL PROCEDURES

### Mice

All animal studies conformed to guidelines set by the Office of the Institutional Animal Care and Use Committee (IACUC) and performed with IACUC approval. FVB/NJ and KRT5-Cre^ERT2^ mice (FVB.Cg-Tg(KRT5-cre/ER^T2^)2Ipc/JeldJ) mice were purchased from Jackson Laboratories (stock no 001800 and 018394). Tfap2c^fl/fl^ mouse strain was previously described (Auman et al., 2002). All mice used in experiments were backcrossed >15 generations on FVB/NJ background to limit genetic variability. Knockout mice were generated by crossing *Tfap2c*^fl/fl^ with K5-Cre^ERT2^ mice, and the offspring were back crossed with *Tfap2c*^fl/fl^ to generate FVB/*Tfap2c^fl/fl^/Krt5-* Cre^ERT2^. Pulsing with 4OH-Tamoxifen was performed starting at 4-weeks of age at 30 mg/kg/day intraperitoneal injection x 5 days. Mouse pair 4 mammary glands were used for experimental analysis. Genotype analysis was conducted with qPCR analysis of tail DNA by Transnetyx, Inc. (Cordova, TN). Only female mice were used for analysis.

### Mammary cell preparation, labeling, and flow cytometry

The 4^th^ mammary gland of two corn oil treated and two 4OHT treated 8 to 9-week-old virgin female FVB/*Tfap2c^fl/fl^* mice were digested in a mixture of gentle collagenase/hyaluronidase (Stemcell, Catalog #07919) with 5% fetal bovine serum (FBS) in complete EpiCult™-B (Stemcell, #05611) media for 15 hours at 37 °C. The red blood cells were lysed in NH4Cl. The cell suspension was further digested in 0.25% trypsin-EDTA for 1 minute then 5 U/ml dispase (Stemcell, #07913) plus 0.1 mg/ml DNase I (Stemcell, #07900) for 1 minute followed by filtration through a 40-μm cell strainer. After washing in HBSS, cell viability was assessed by Trypan Blue. Single cell isolation procedure was performed as previously described (Borcherding et al., 2015). Fluorescence-activated cell sorting (FACS) was carried out by FACS Aria (Becton Dickinson). The Lin^−^ population was defined as Ter119^−^, CD31^−^, CD45^−^. FACS data were analyzed using FlowJo software (v 10.1r7, Tree Star). The following antibodies were used: CD31-Biotin (Biolegend, 102404, 1:250), TER119-Biotin (Biolegend, 116204, 1:250), CD45-Biotin (Biolegend, 103014, 1:250), Strep-PE-Cy7 (Invitrogen, 25-4317-82, 1:250), CD24-PE (Biolegend, 101807, 1:250), CD49f-FITC (Biolegend, 313606, 1:200), Procr-APC (Biolegend, 141505, 1:250) for staining and Procr-APC (Invitrogen, 25-5931-82) for compensation with compensation beads (Invitrogen 01-2222-42).

### RNA Analysis

RNA was isolated from basal and luminal cells recovered from flow cytometry using the RNeasy Mini Kit (Qiagen, catalog no. 74104). Using the random hexamers method (Thermo Fisher Scientific), mRNA was converted to cDNA by quantitative PCR (qPCR). Relative expression of exon 2/3 of *Tfap2c* (TFS, catalog no. 4351372, ID: Mm00493470_g1) and exon 6 of *Tfap2c* (TFS, catalog no. 4351372, ID: Mm00493474_m1) was determined using TaqMan primers and the ΔΔ*C*t method of qPCR. Amplification of *Gapdh* (TFS, catalog no. 4331182, ID: Mm99999915_g1) was used as an endogenous control.

### Mammosphere assay

For mammosphere assays, freshly sorted cells were cultured on low attachment plates (Corning, #3473) in EpiCult™-B Basal Medium (Stemcell, #05611) supplemented with EpiCult™-B Proliferation Supplement (#05612), 10 ng/mL epidermal growth factor, 10 ng/mL βFGF, 4 μg/mL, 10% FBS, and 50 μg/mL gentamicin (IBI Scientific, #IB02030) in a low-oxygen incubator at 37 °C for 21 days (Shaw et al., 2012). At 21 days, secondary mammospheres were generated from introduction of trypsin and mechanical disruption of primary mammospheres. Secondary mammospheres were incubated in low-oxygen incubator at 37 °C for 14 days. Images for comparative analysis were taken at 14 days. Mammosphere count was evaluated by ImageJ using 50 μm minimum diameter and evidence of 3-dimensional structure to differential from cellular aggregates.

### Single cell capture and library preparation for sequencing

Freshly sorted epithelial cells were submitted for sequencing using the manufacturer recommended Chromium (10x Genomics) and Illumina technologies. The amplified cDNA was constructed into 3’ expression libraries and pooled together in separate lanes of a 150 base-pair, paired-end chemistry flow cell. The Illumina HiSeq 4000 was used with Agilent Bioanalyzer, and average library sizes ranged from 329 bp to 432 bp in length. The adapter sequence of the library was 124 bases in length. Basecalls were converted into FASTQ files by the University of Iowa Genomics Division using the Illumina bcl2fastq software. These samples captured an estimated 3,546 to 5,545 number of cells with mean reads per cell ranging from 63,842 to 94,776. Data is available in GEO under the accession GSE143159.

### scRNA-seq data processing and quality control

Transcripts were mapped using Cell Ranger Version 3.0.1 and mm10 reference genome (GRCm38.91). Using R, the filtered gene count matrix was imported, and quality control for all samples was completed prior to downstream analysis using the standard workflow put forth by the Seurat R package, version 3.1(Butler et al., 2018; Stuart et al., 2019). All cells with unique RNA feature counts less than 200, or the percent mitochondrial DNA > 5% were removed.

### scRNA-seq clustering and visualization

scRNA-seq data normalization using Seurat, was completed to account for feature expression measurements in each cell in relation to total expression values and the data were log-transformed using a 1e4 factor(Butler et al., 2018; Stuart et al., 2019). The subset of features showing high cell-to-cell variability were identified in the dataset. The individual runs were combined using mutual nearest neighbor approach as previously described(Stuart et al., 2019), enabling further analysis to allow for identification of common cell types between datasets and to control for batch effect between samples. Linear scaling was performed and principal component analysis (PCA) calculations were completed. Uniform Manifold Approximation and Projection (UMAP) was calculated using the *RunUmap* function with default settings and PC input equal to 20(Becht et al., 2018). The *FindClusters* with default Louvain algorithm and a resolution of 0.5 clustered cells was completed, identifing 13 clusters. Differential gene expression was calculated using the Wilcoxon rank sum test with a pseudocount of 0.01 and without thresholds for log-fold change or percentage of cells. P-value was adjusted using the Bonferroni method for multiple hypotheses comparison.

### Pathway Analysis

Differential gene expression results derived from the scRNA-seq quantifications were analyzed using Ingenuity Pathway Analysis (IPA, QIAGEN Inc., Hilden, Germany). Each cluster was individually analyzed to examine the relationship between its highly significantly up- or downregulated genes between CO versus 4OHT. Analysis was completed with adjusted p value < 0.05 and log-fold change between 0.5-2.0 in order to isolate the top 3-5 functions altered in each cluster. Functions of most interest were related to mammary gland development such as cell movement, cell development, tissue morphology, and connective tissue development and function.

### Cell cycle analysis

To determine cell cycle heterogeneity, cell cycle phase was calculated with scRNA-seq data in both CO and 4OHT treated mice on the previously generated integrated object. Scores were determined based on previously published markers.(Nestorowa et al., 2016) Utilizing the *CellCycleScoring* function in Seurat on integrated RNA, S and G2/M scores were stored in the object meta data along with classification of each cell as G2M, S, and G1 phase. Cell cycle distribution based on cluster was visualized via creating a frequency table and graphs using the ggplot2 R package, version 3.2.1.

### Pseudotime analysis

Using Monocle2, pseudotime analysis was completed on the corn oil data subset of the previously generated scRNA-seq Seurat integrated object(Qiu et al., 2017a; Qiu et al., 2017b; Trapnell et al., 2014). Cells were reclustered using the *reducedDimension* function in monocle that utilized five dimensions to generate a t-distributed stochastic neighbor embedding (tSNE) projection. Differential genes by the monocle clusters were calculated by vector generalized linear and additive modelling with isolating the top 1,000 genes with the lowest q value as ordering genes. Dimension reduction was completed using the DDRtree method with the residual model based on the single-cell run. Cluster numbers were the same as previously assigned numbers from Seurat analysis. The branch with the majority of Procr+ stem cells was selected as the pseudotime root state.

### Adenovirus infection

Ad5CMVeGFP and Ad5CMVCre-eGFP were purchased from the University of Iowa DNA Core Laboratory. Virus infection was performed at a multiplicity of infection (MOI) of 200. All experiments were performed with cells harvested from cell sorting.

### Mouse mammary gland transplantation

3-week-old FVB/NJ recipient females were anesthetized, and the fat pad of right 4^th^ mammary gland was cleared up to the lymph node. In one set of experiments, MMECs harvested from FVB/*Tfap2c^fl/fl^* mouse mammary glands were infected with adenovirus and transplanted into the cleared fat pad of recipient mice. In other experiments, approximately 1 mm^3^ sections of mammary gland were harvested from 9-week old FVB/Tfap2cfl/fl/Krt5-Cre-ERT2 mice that had been treated with 4OHT or corn oil from 4-to 5-weeks and were transplanted into the cleared fat pads. After eight weeks, transplanted mammary glands were collected for whole-mount staining and analysis.

### Whole-mount analysis of mammary branching

Mammary glands were collected and whole-mounts performed as previously described (Plante et al., 2011). In brief, the 4^th^ mammary glands were fixed in Carnoy’s fixative solution at 4 °C overnight. After washing in a series of ethanol dilutions, sections were stained with the Carmine Alum solution at room temperature overnight. The following morning the mammary glands were washed in a series of ethanol dilutions and cleared in Histo-Clear overnight. To analyze branching morphogenesis, whole-mount mammary gland images were taken on a light box, and branching distance was measured from lymph node to the furthest terminal end buds. For mammary gland transplantation experiments, the branching area was measured. All images were quantified by ImageJ.

### Immunohistochemistry

Immunohistochemical analysis was accomplished on formalin-fixed, paraffin-embedded samples from the 4^th^ mammary gland. The following antibodies were used for analysis: anti-AP-2γ, 1:100 (Santa Cruz Company, #sc-12762); anti-CRE, 1:50 (Cell Signaling, #15036); anti-CK5, 1:100 (Abcam, ab5235) and anti-CK8/18, 1:500 (Abcam, ab53280). Standard protocols were followed, and Vector ABC kits used for amplification. Brightfield imaging was collected on an Olympus BX-51 microscope.

### Statistical Analysis

For statistical analyses of whole-mount and immunohistochemical data, Mann-Whitney U test and Fisher’s exact test were performed using GraphPad Prism; P-values of less than 0.05 were considered significant. Mammosphere counts and sizes were analyzed using chi-square test. Proportion z-test was used to determine significance between scRNA-seq cluster cell counts from Seurat.

## AUTHOR CONTRIBUTIONS

Conceptualization: VWG, EC, TW, WZ, RJW; Methodology: VWG, EC, DTT, TW, WZ, RJW; Software: EC, DTT, NB, MVK; Validation: VWG, EC, DTT, VCC, NB, KEK, VTW, AWL, MVK; Formal Analysis: VWG, EC, DTT, VCC, NB, VTW, AWL, MVK; Investigation: VWG, EC, DTT, VCC, KEK, VTW, AWL; Resources: VWG, EC, DTT, KEK; Data Curation: EC, DTT, NB, MVK; Writing—Original Draft: VWG, RJW; Writing— Review & Editing: VWG, EC, DTT, VCC, NB, KEK, VTW, AWL, MVK, TW, WZ, RJW; Visualization: VWG, EC, DTT, KEK; Supervision: VWG, WZ, RJW; Project Administration: WZ, RJW; Funding Acquisition: TW, WZ, RJW

## ACKNOWLEDGMENTS

This work was supported by the NIH grants R01CA183702 (PI: R.J.W.), CA200673 (PI: W.Z.), CA203834 (PI: W.Z.) T32CA148062 (P.I.: R.J.W.), DOD/CDMRP grant BC180227 (PI: W.Z.) and by a generous gift from the Kristen Olewine Milke Breast Cancer Research Fund (to R.J.W.) and a gift from Dr. and Mrs. James Robert Spencer Family Cancer Research Fund (to W.Z.). E.C., D.T.T., K.E.K., V.T.W. and A.W.L. were supported by the NIH grant T32CA148062. N.B. was supported by a F30 fellowship CA206255.

