## Supplemental 1-7 for "AP-2γ is Required for Maintenance of Pluripotent Mammary Stem Cells"

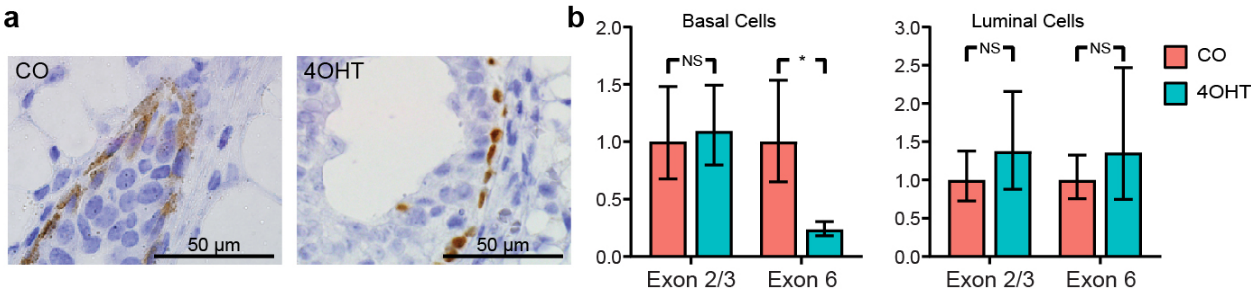


**Figure S1. 4OHT induces nuclear Cre and excision of Tfap2c exon 6. a.** FVB/*Tfap2c*$^{fl/fl}$/*Krt5-Cre-ER*$^{T2}$ mice were pulsed with corn oil (CO) or 4-hydroxy-tamoxifen (4OHT) and Cre expression was examined by IHC; treatment with 4OHT induced nuclear Cre expression. **b.** *Tfap2c* RNA from FVB/*Tfap2c*$^{fl/fl}$/*Krt5-Cre-ER*$^{T2}$ mice treated with CO or 4OHT was analyzed by RT-PCR using primers that amplify across exons 2/3 or exon 6 of the *Tfap2c* gene; data demonstrates loss of *Tfap2c* RNA containing exon 6 in basal cells recovered from mice treated with 4OHT.

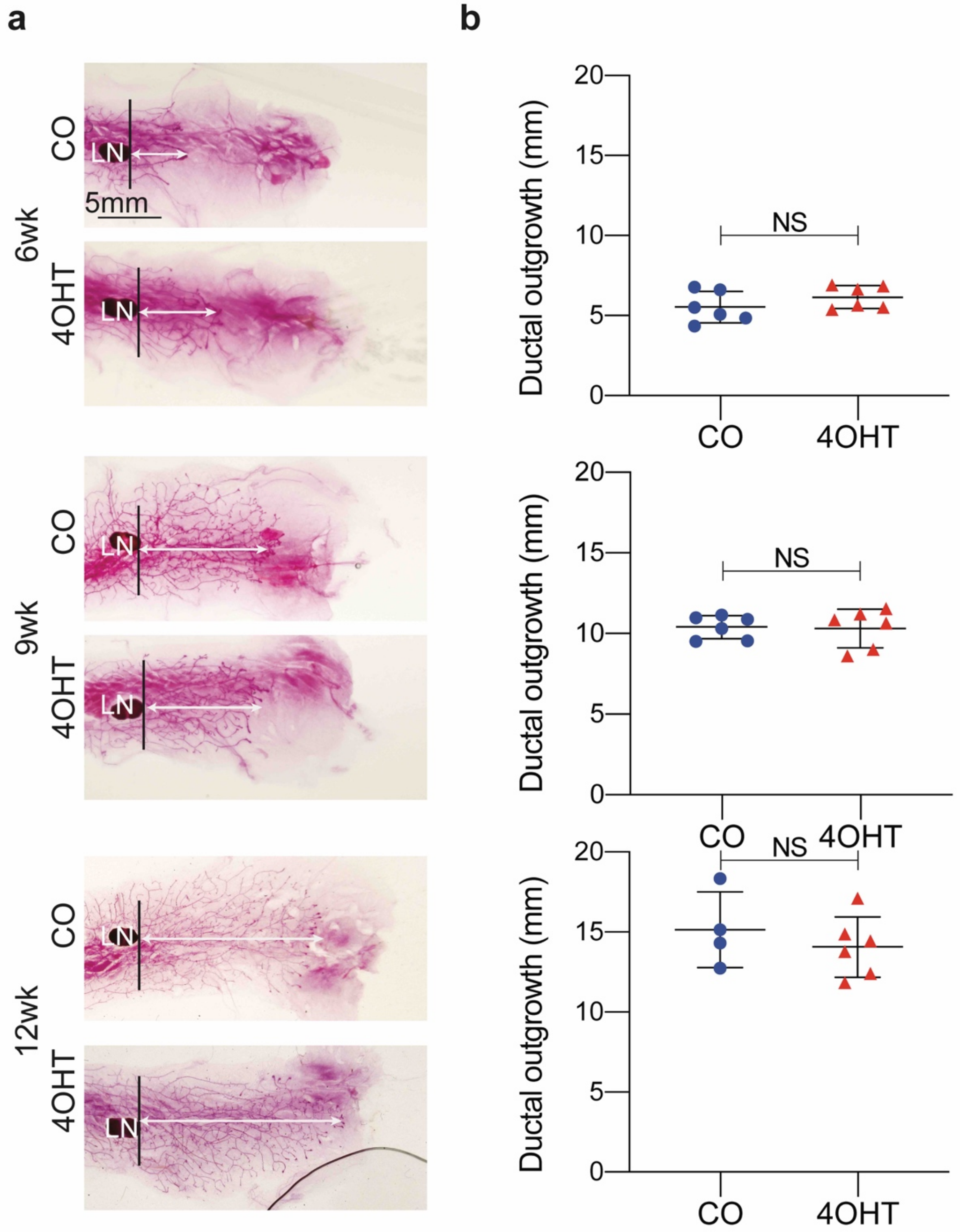


**Figure S2. 4OHT does not alter mammary gland ductal outgrowth. a.** Examples of whole mounts from FVB/*Krt5-Cre-ER*$^{T2}$ mice treated with CO or 4OHT for five days starting at 4 weeks of age and analyzed for ductal development at 6, 9 and 12 weeks. **b.** Analysis of ductal outgrowth shows no differences for mice treated with CO vs. 4OHT.

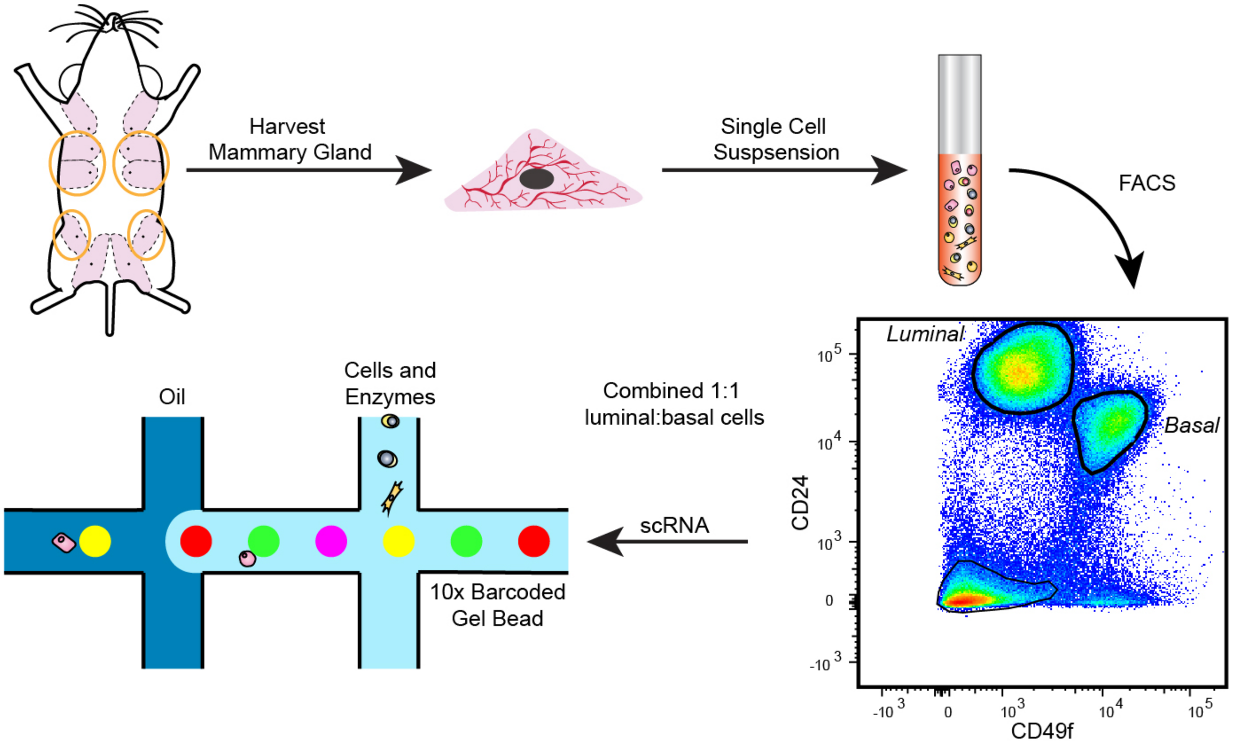


**Figure S3. Diagram of scRNA-seq analysis.** The 4$^{th}$ mammary glands of 9-week-old virgin female FVB/*Tfap2c*$^{fl/fl}$ mice, which had been treated with CO or 4OHT at 4 weeks of age, were harvested and single cell suspensions created; FACS analysis was performed and the luminal and basal MMECs were recovered; luminal and basal cells were mixed 1:1 and subjected to RNA-seq.

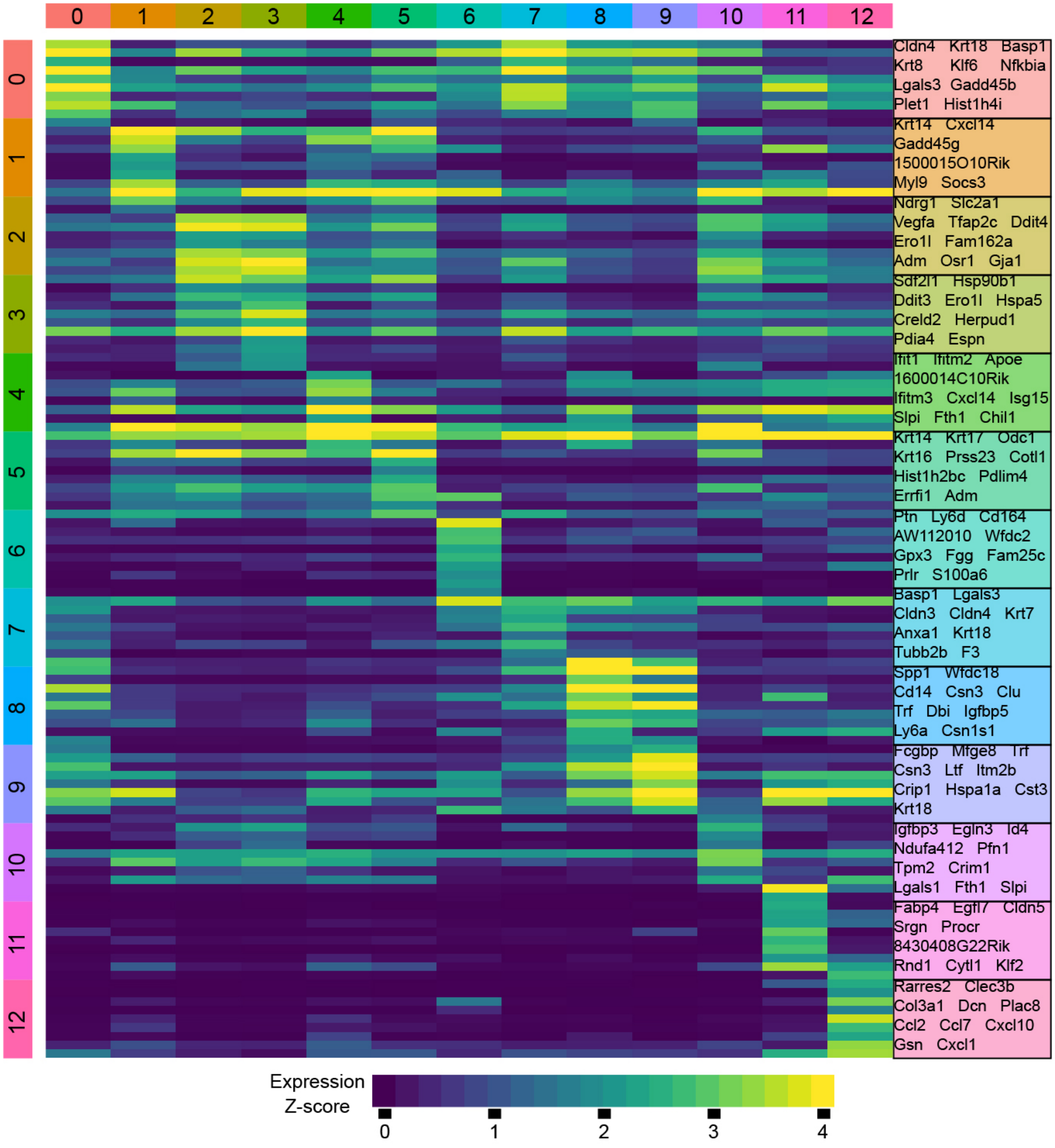


**Figure S4. Transcriptome of MMECs clusters.** Transcriptome of the 13 MMECs clusters showing the top 10 genes characteristic of each cluster.

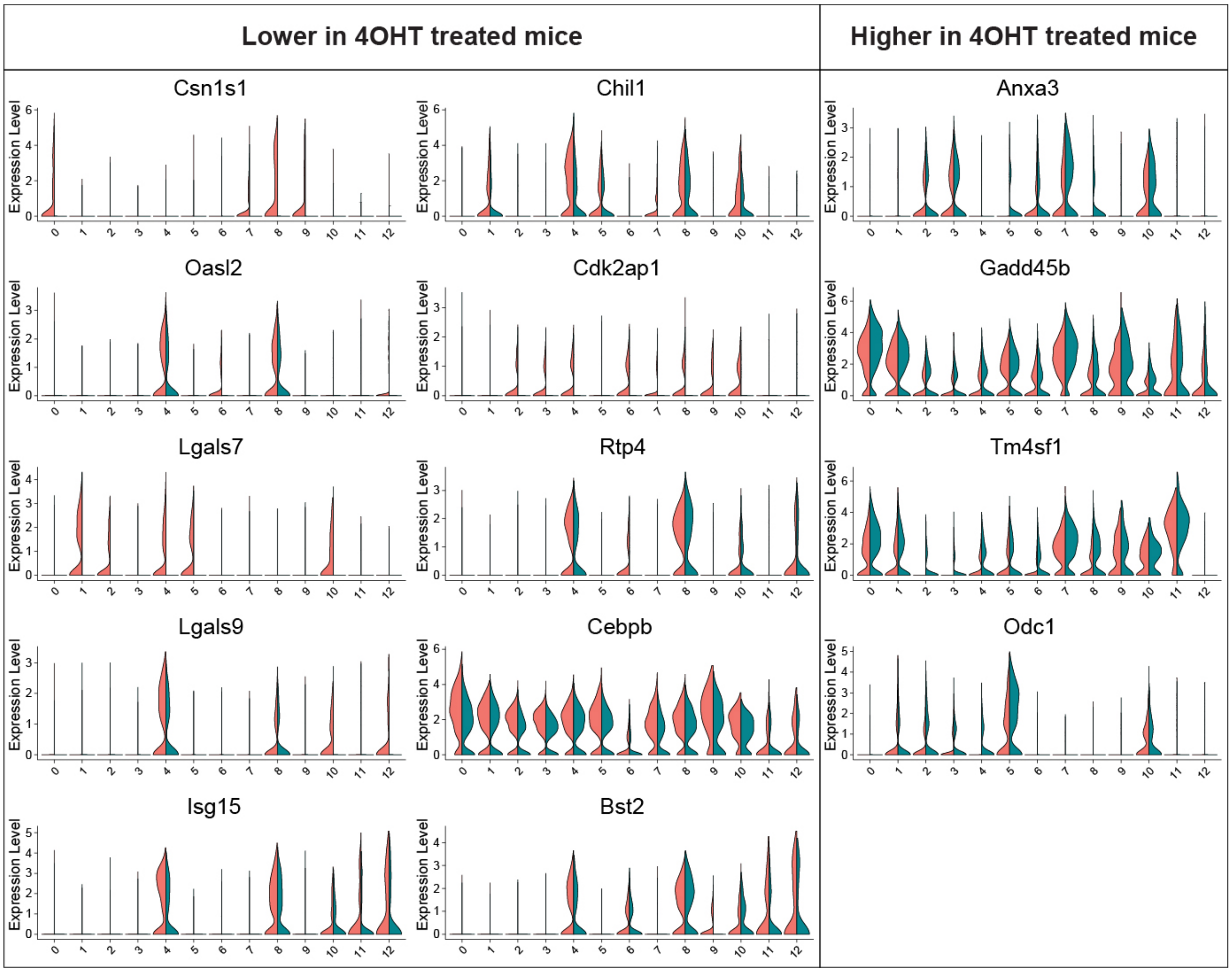


**Figure S5. Violin plots of genes altered by *Tfap2c* CKO by MMEC cluster.** Violin plots showing details of expression for top 10 genes lower with CKO of *Tfap2c* (left panels) and top four genes higher with CKO of *Tfap2c* (right panels) by cluster.

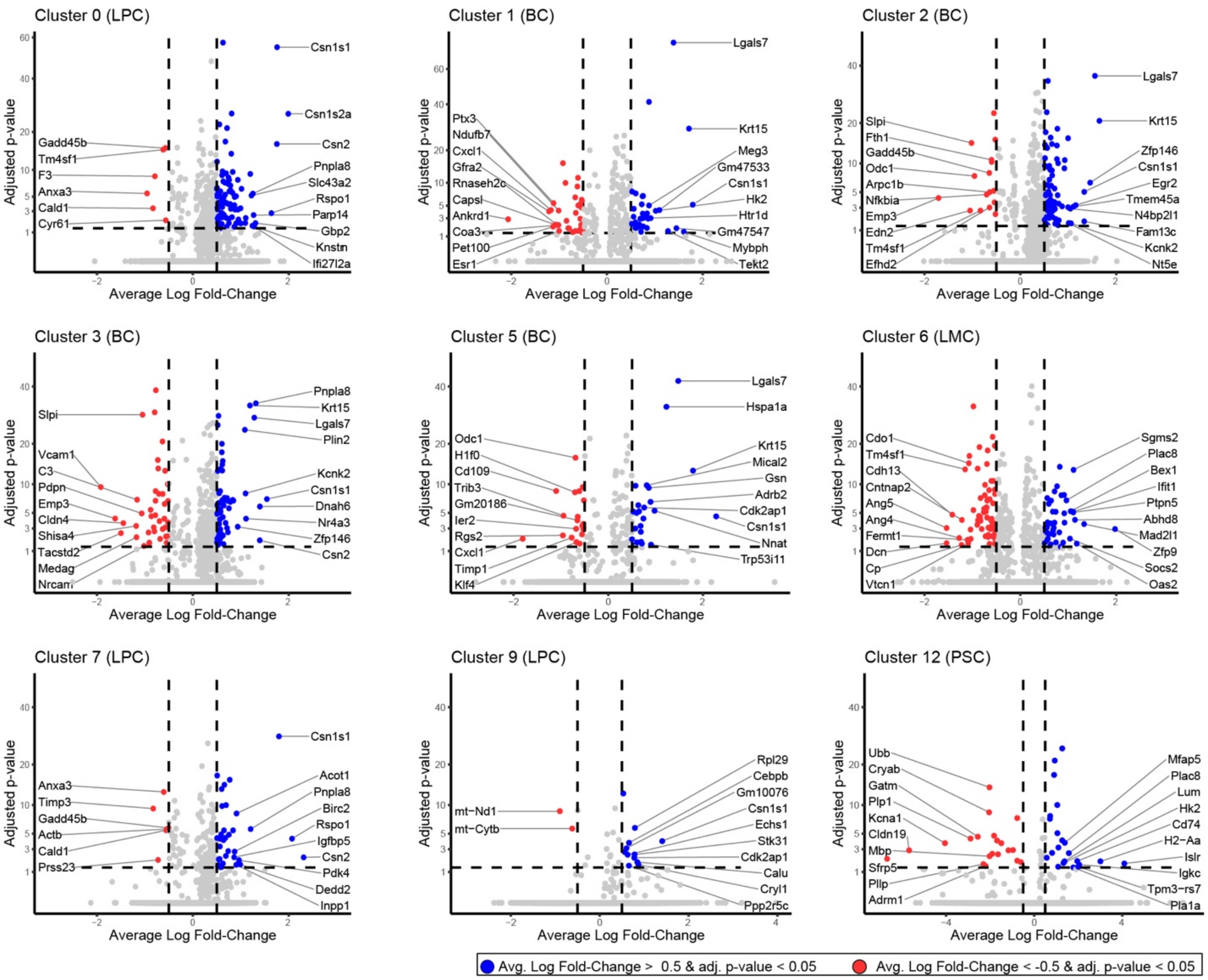


**Figure S6. Volcano plots for remaining MMEC clusters.** Volcano plots showing changes in gene expression with CKO of *Tfap2c* for the remaining clusters not included in Figure 4.

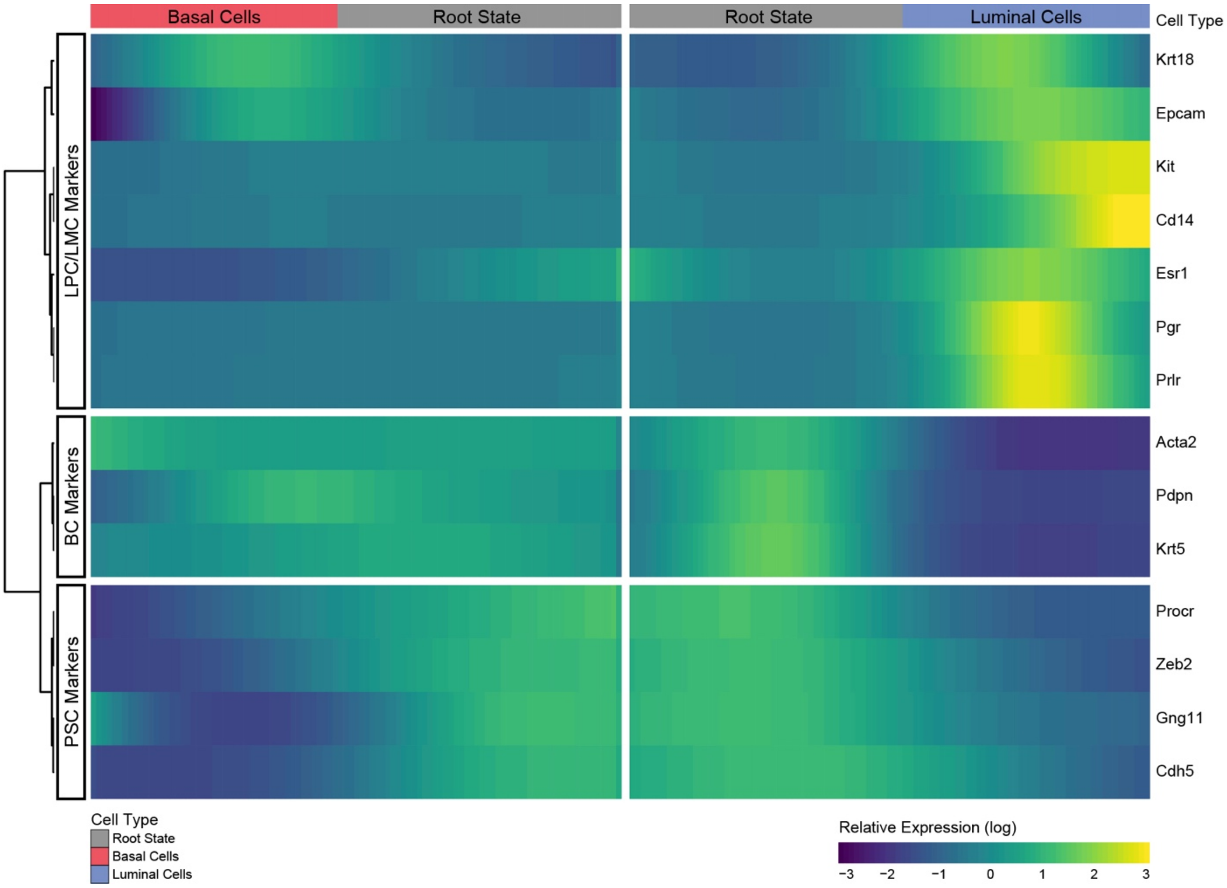


**Figure S7. Heatmap from Pseudotime analysis.** Heatmap showing examples of gene expression for Pseudotime analysis demonstrating progression of gene expression from the root state with MaSC markers to the basal and luminal patterns of expression.
